# The RNA binding protein HNRNPA2B1 regulates RNA abundance and motor protein activity in neurites

**DOI:** 10.1101/2024.08.26.609768

**Authors:** Joelle Lo, Katherine F. Vaeth, Gurprit Bhardwaj, Neelanjan Mukherjee, Holger A. Russ, Jeffrey K. Moore, J. Matthew Taliaferro

## Abstract

RNA molecules are localized to subcellular regions through interactions between localization-regulatory cis-elements and trans-acting RNA binding proteins (RBPs). However, the identities of RNAs whose localization is regulated by a specific RBP as well as the impacts of that RNA localization on cell function have generally remained unknown. Here, we demonstrate that the RBP HNRNPA2B1 acts to keep specific RNAs out of neuronal projections. Using subcellular fractionation, high-throughput sequencing, and single molecule RNA FISH, we find that hundreds of RNAs demonstrate markedly increased abundance in neurites in HNRNPA2B1 knockout cells. These RNAs often encode motor proteins and are enriched for known HNRNPA2B1 binding sites and motifs in their 3′ UTRs. The speed and processivity of microtubule-based transport is impaired in these cells, specifically in their neurites. HNRNPA2B1 point mutations that increase its cytoplasmic abundance relative to wildtype lead to stronger suppression of RNA mislocalization defects than seen with wildtype HNRNPA2B1. We further find that the subcellular localizations of HNRNPA2B1 target RNAs are sensitive to perturbations of RNA decay machinery, suggesting that it is HNRNPA2B1’s known role in regulating RNA stability that may explain these observations. These findings establish HNRNPA2B1 as a negative regulator of neurite RNA abundance and link the subcellular activities of motor proteins with the subcellular abundance of the RNAs that encode them.

## INTRODUCTION

The localization of RNAs to specific subcellular locations allows the spatial and temporal control of gene expression. Thousands of RNAs are known to be asymmetrically localized within cells and subject to local regulation (Cajigas et al. 2012; Zappulo et al. 2017; Taliaferro et al. 2016; Engel et al. 2020; Adekunle and Wang 2020; Moor et al. 2017; Lécuyer et al. 2007). The localization of some of these RNAs is critical for cell function and organismal development in species ranging from yeast to Drosophila to mammals (Long et al. 1997; Ephrussi et al. 1991; Lécuyer et al. 2007; Kislauskis et al. 1994; Eom et al. 2003).

For a handful of well-studied RNAs, the mechanisms that underlie their transport is well understood (Bertrand et al. 1998; Ephrussi et al. 1991; Forrest and Gavis 2003; Safieddine et al. 2021; Goering et al. 2023; Engel et al. 2020; Arora et al. 2022a; Kislauskis et al. 1994). Generally, these RNAs contain localization-regulatory cis-elements within them, often in their 3′ UTRs, that are bound by RBPs that mediate transport. These elements were identified through careful yet laborious experiments that often involved repeated mutation and/or deletion of sequences within their 3′ UTRs in order to identify functional elements.

However, for the vast majority of localized RNAs, the sequences within them and the identities of the RBPs that regulate their localization are unknown. Newer techniques for identifying RNA localization mechanisms have moved from the single gene-focused approaches of earlier studies to the transcriptome-wide quantification of RNA localization (Taliaferro 2022; Arora et al. 2021; Fazal et al. 2019; Padrón et al. 2019; Engel et al. 2021). By identifying sets of transcripts whose localization depends upon a given RBP, these approaches allow the identification of commonalities shared among the localization-regulatory targets of the RBP. These commonalities often shed light on general rules that define sequence features that regulate RNA localization as well as the RBPs that bind them.

The *Hnrnpa2b1* gene produces two protein isoforms, HNRNPA2 and HNRNPB1. These isoforms differ only at their N-terminus with HNRNPB1 containing an additional 12 amino acids. Little is known about if and how these highly related isoforms behave differently (Nguyen et al. 2018), and they are often referred to together as HNRNPA2B1. HNRNPA2B1 regulates a range of post-transcriptional regulatory processes, including splicing, 3′ end formation, and RNA decay (Martinez et al. 2016; Huelga et al. 2012; Liu and Shi 2021), and mutations within HNRNPA2B1 that affect its ability to perform these actions are associated with neurodegenerative diseases (Martinez et al. 2016). Across these studies, HNRNPA2B1 has been found to bind a range of RNA sequences but has been repeatedly found to bind to sequences rich in adenosine and guanosine (Martinez et al. 2016; Huelga et al. 2012; Dominguez et al. 2018; Kasim et al. 2014). Although some evidence exists that HNRNPA2B1 regulates RNA localization (Ainger et al. 1997; Shan et al. 2000), the set of transcripts that depend upon HNRNPA2B1 for proper localization as well as the phenotypic consequences of their mislocalization are currently unknown. In this study, we address that knowledge gap by profiling subcellular transcriptomes from HNRNPA2B1 knockout and rescue neuronal cells.

## RESULTS

### Generation of HNRNPA2B1 knockout and rescue cells

We and others have previously used compartmentalized culture systems that allow the separation of neuronal cells into soma and neurite fractions (Taliaferro et al. 2016; Goering et al. 2020, 2023; Arora et al. 2021; Zappulo et al. 2017; Ciolli Mattioli et al. 2019). These systems work by culturing cells on porous membranes. The pores in the membranes allow neurite growth to the underside of the membrane but restrict cell bodies to the top of the membrane. Cells can then be mechanically fractionated into soma and neurite fractions by scraping the top of the membrane. We then collect RNA from each fraction and analyze it by high-throughput sequencing to determine relative neurite enrichments for all cellular RNAs (**Figure 1A**).

**Figure 1.**
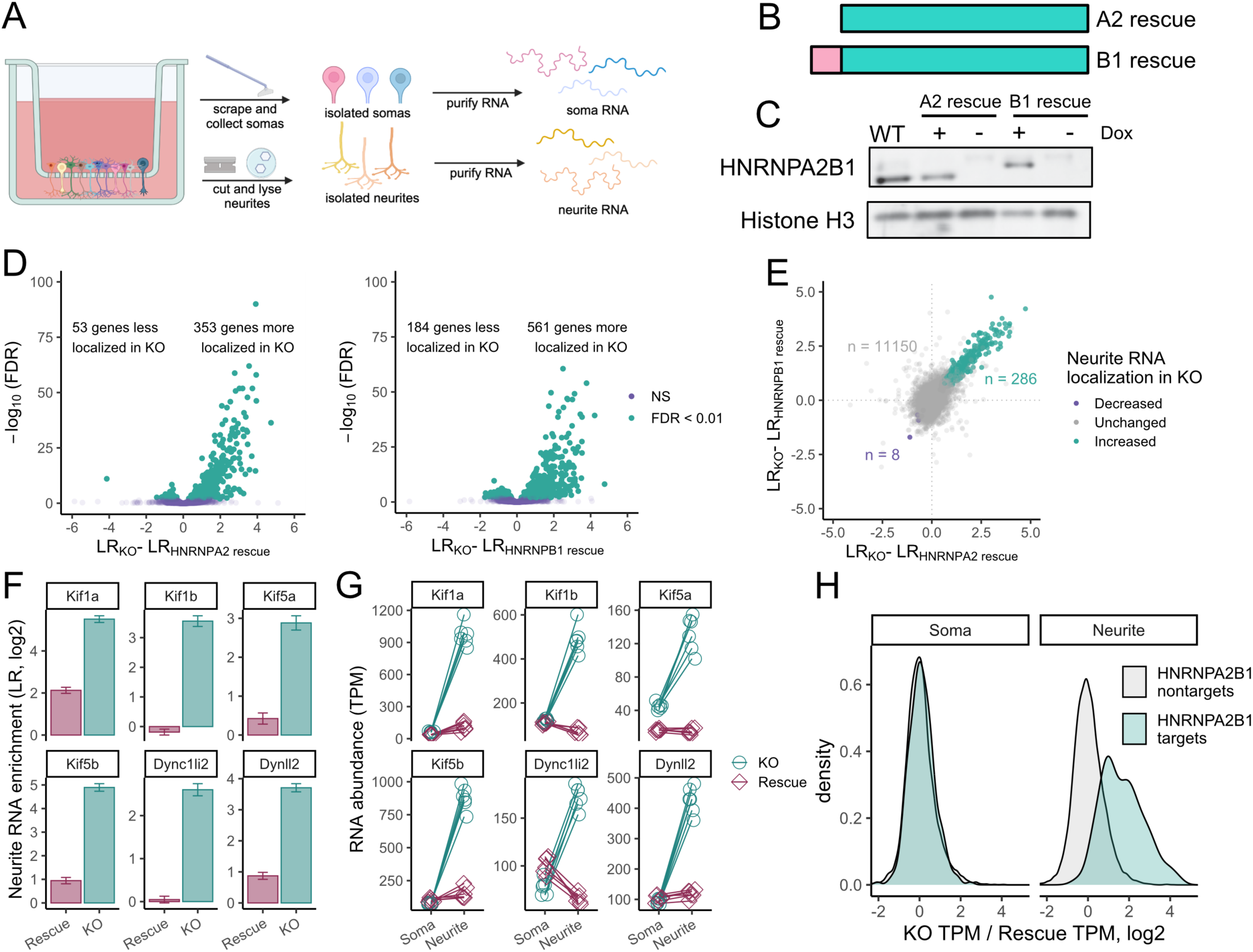
HNRNPA2B1 regulates the localization of hundreds of RNAs. (A) Schematic of subcellular fractionation. RNA samples are collected from soma and neurite fractions and profiled using high-throughput sequencing. (B) Schematic of HNRNPA2 and HNRNPB1 protein isoforms. (C) Immunoblot of doxycycline-inducible HNRNPA2 and HNRNPB1 rescue cells. For comparison, lysate from wildtype, unmodified cells is also shown in which the HNRNPA2 is predominantly expressed. (D) Changes in LR values comparing knockout and HNRNPA2 (left) or HNRNPB1 (right) rescue cells. (E) Comparison of RNA localization results from HNRNPA2 and HNRNPB1 rescue cells. Genes marked as displaying increased (teal) or decreased (purple) neurite-enrichments were identified as such in both HNRNPA2 and HNRNPB1 experiments (FDR < 0.01). (F) Changes in neurite enrichments between HNRNPA2B1 knockout and rescue cells for selected RNAs that encode motor proteins. (G) Changes in soma and neurite abundances between HNRNPA2B1 knockout and rescue cells for selected RNAs that encode motor proteins. (H) Changes in soma and neurite abundances between HNRNPA2B1 knockout and rescue cells for RNAs identified as HNRNPA2B1 localization targets (teal) or HNRNPA2B1 localization-insensitive (gray) in figure 1E.

We set out to use this technique to determine which RNAs relied on HNRNPA2B1 for proper localization in neuronal cells. To facilitate this, we created an HNRNPA2B1 knockout CAD (Cath.a-differentiated) cell line. CAD cells are derived from a mouse brain tumor and can be induced through serum starvation to display a number of neuronal characteristics, including the upregulation of neuronal markers, the downregulation of the cell cycle, and a distinct neuronal morphology (Qi et al. 1997). Using CRISPR/Cas9 and guide RNAs targeted to the 5′ and 3′ ends of the *Hnrnpa2b1* gene, we removed the entire locus.

The *Hnrnpa2b1* locus expresses two distinct yet highly related protein isoforms, HNRNPA2 and HNRNPB1. These isoforms differ only in their N-termini with the HNRNPB1 isoform containing an extra 12 amino acids (**Figure 1B**). We wanted to be able to separately test the ability of each isoform to regulate RNA localization. Using *Cre*-mediated site-specific recombination, we rescued these knockout cells by integrating transgenes encoding either HNRNPA2 or HNRNPB1. The expression of these transgenes was under the control of doxycycline, allowing us to assay knockout and rescue conditions by omitting and adding doxycycline, respectively (**Figure 1C**). Using immunoblotting, we found that the expression of each rescuing transgene was approximately equal to the expression from the *Hnrnpa2b1* locus observed in wildtype cells (**Figure 1C**).

### HNRNPA2 and HNRNPB1 regulate the localization of hundreds of RNAs

To identify RNAs whose localization was sensitive to the loss of HNRNPA2 and/or HNRNPB1, we cultured HNRNPA2 and HNRNPB1 rescue cells on porous membranes in the absence (knockout) and presence (rescue) of doxycycline. We then isolated RNA from soma and neurite fractions of the cells and quantified it using RNAseq. Principal component analysis of the RNAseq data indicated that samples separated by subcellular compartment and genotype along the first two principal components (**Figures S1A-S1C**), and hierarchical clustering analysis revealed that samples separated first by genotype and then by subcellular compartment (**Figure S1D**).

For each RNA, we used a metric called “Localization Ratio” (LR) to quantify its enrichment in the neurite fraction compared to the soma fraction. The LR for each RNA is defined as the log_2_ ratio of its relative abundance in the neurite compared to its relative abundance in the soma. RNAs that are enriched in the neurite therefore have positive LR values while those that are enriched in the soma have negative LR values.

To validate the efficiency of the subcellular fractionation, we compared LR values for mRNAs encoding ribosomal proteins and components of the electron transport chain to LR values for all other mRNAs. We and others have previously found these two classes of RNAs to be strongly and consistently neurite-enriched (Taliaferro et al. 2016; Goering et al. 2020; Zappulo et al. 2017). They therefore serve as positive controls for a successful separation of soma and neurite RNA. Encouragingly, we found that in all samples, RNAs from these two classes had significantly higher LR values than other RNAs, indicating that the subcellular fractionation and RNA isolation were successful (**Figure S1E**).

By comparing LR values in knockout and rescue cells, we identified RNAs whose localization was dependent on HNRNPA2 and/or HNRNPB1. Interestingly, we found that many RNAs were *more* neurite enriched in knockout cells compared to those rescued with either HNRNPA2 or HNRNPB1, indicating that these proteins were acting to keep hundreds of RNAs out of neurites (**Figure 1D, Table S1, Table S2**). To ask if the two protein isoforms regulated the same or different RNA targets, we compared changes in LR values observed between knockout and HNRNPA2 rescue cells to those observed between knockout and HNRNPB1 rescue cells. We found a strong correlation between changes in LR values induced by the two rescue isoforms, indicating that both isoforms were regulating the localization of largely the same set of RNAs (**Figure 1E**). We therefore chose to move forward with 286 RNAs that were significantly more neurite-enriched upon the loss of both HNRNPA2 and HNRNPB1 for further analysis (FDR < 0.01, **Figure 1E**). We will refer to these RNAs as HNRNPA2B1-target RNAs. Since the rescue isoforms regulated largely the same RNAs, subsequent experiments were done using only knockout and HNRNPA2-rescue samples. For simplicity, we will still refer to the protein as HNRNPA2B1, even though only the HNRNPA2 isoform will be used in further experiments. However, given the above findings, the results will likely be applicable to both HNRNPA2 and HNRNPB1 isoforms.

### Several RNAs encoding motor proteins display HNRNPA2B1-sensitive localization patterns

We analyzed the identities of the HNRNPA2B1-target RNAs using gene ontology analysis (Eden et al. 2009). We found that most of the top enriched terms dealt with subcellular transport (**Figure S1F**). Indeed, RNAs encoding several components of motor proteins, including both kinesins and dyneins, were strongly more neurite enriched in knockout cells than in HNRNPA2B1 rescue cells (**Figure 1F**).

In principle, because the LR metric is a ratio, the increase in LR value observed in knockout cells could be due to either an increase in the numerator of the ratio (neurite abundance) or a decrease in the denominator (soma abundance) or both. To answer this, we compared RNA abundances in soma and neurites separately for several affected genes. We observed that when comparing knockout and rescue samples, soma RNA abundances for these genes were generally unchanged. However, their abundances in neurites were greatly increased in knockout cells by up to 20-fold (**Figure 1G**). This effect extended beyond a small handful of RNAs and could be observed throughout the 286 HNRNPA2B1-target RNAs (**Figure 1H**).

These results indicate that we are observing a neurite-specific increase in the levels of the mislocalized RNAs. Further, because the soma contains approximately 95% of the cellular RNA and abundances in this fraction are generally unchanged, we are not observing a broad, cell-wide upregulation of these RNAs.

### RNA mislocalization in HNRNPA2B1 knockout cells is validated by smFISH

To validate RNA mislocalization patterns we observed using compartment-specific RNAseq, we used single molecule RNA FISH (smFISH) to probe the localization patterns of three motor protein mRNAs in HNRNPA2B1 knockout and rescue cells (Tsanov et al. 2016). We quantified individual puncta corresponding to single RNA molecules for *Kif1a*, *Kif5b*, and *Dynll2* using FISH-quant (Mueller et al. 2013; Arora et al. 2022b) (**Figure 2A-C**). We then counted the number of RNA molecules for each gene in the soma and neurite compartments of differentiated CAD cells. We found that the neurites of HNRNPA2B1 knockout cells contained significantly more RNA molecules encoding *Kif1a*, *Kif5b*, and *Dynll2*, consistent with the subcellular fractionation and RNAseq results (**Figure 2D**).

**Figure 2.**
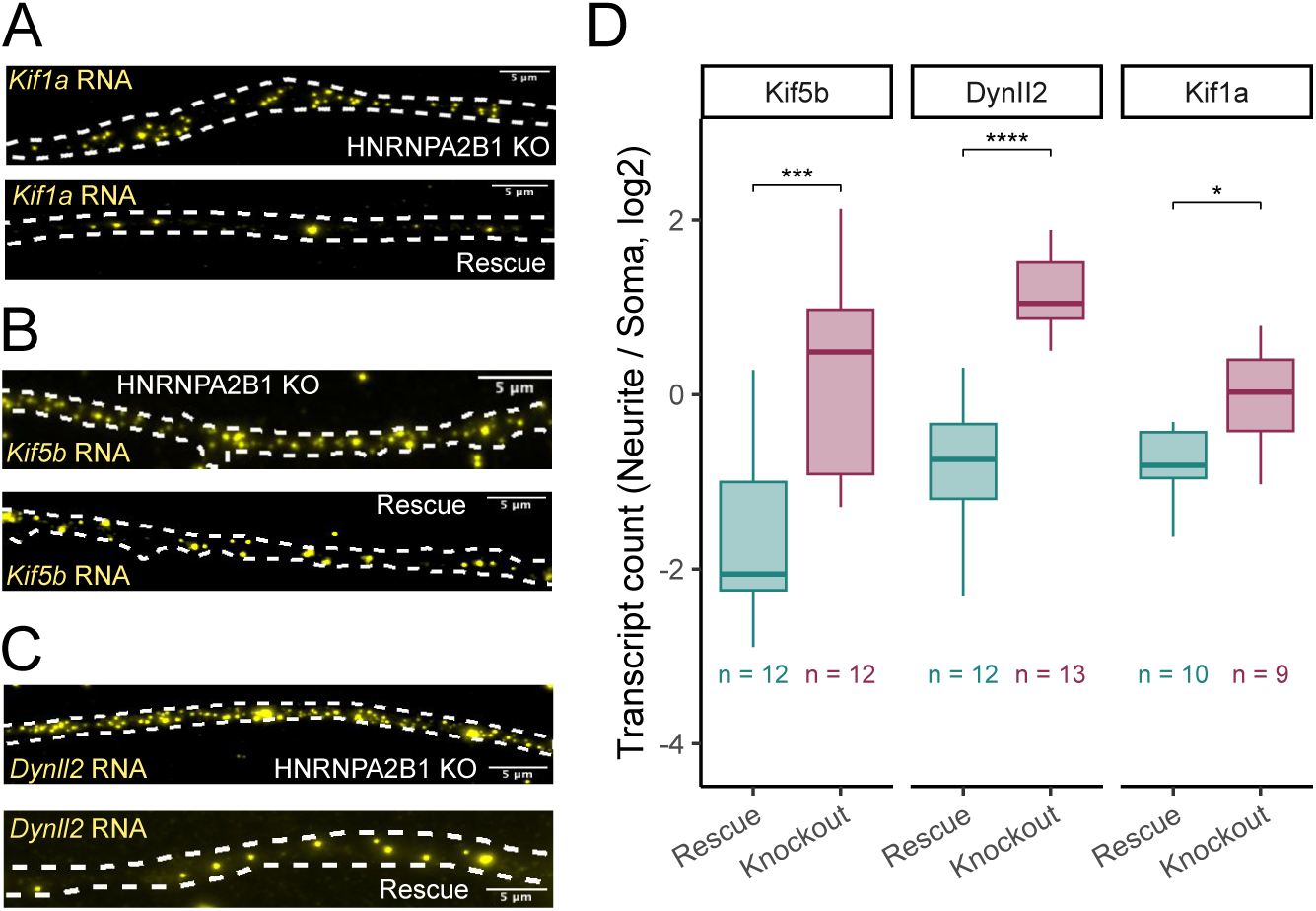
Validation of HNRNPA2B1-dependent RNA localization using smFISH. (A-C) smFISH visualization of Kif1a (A), Kif5b (B), and Dynll2 (C) RNA in neurites of HNRNPA2B1 knockout and rescue cells. Smaller puncta are RNA molecules. Large, bright blotches are aggregates of fluor. These are ignored on the basis of their size and intensity during computational spot calling procedures. (D) Quantitation of smFISH results. RNA molecules are counted in both soma and neurites and compared across conditions. P-values were calculated using a t-test. NS (not significant) represents p > 0.05, * p < 0.05, ** p < 0.01, *** p < 0.001, and **** represents p < 0.0001.

### RNAs with HNRNPA2B1-sensitive localization are enriched for HNRNPA2B1 binding motifs and CLIP sites

If the observed RNA mislocalization patterns of the HNRNPA2B1-target RNAs were due to a loss of direct binding of HNRNPA2B1 to these RNAs, we would expect that the mislocalized RNAs would be enriched for sequences known to be bound by HNRNPA2B1. HNRNPA2B1 has been documented to bind a range of sequence motifs (Liu and Shi 2021), but has consistently been found to prefer sequences rich in adenosine and guanosine (Martinez et al. 2016; Huelga et al. 2012; Muslimov et al. 2014; Munro et al. 1999; Ainger et al. 1997; Dominguez et al. 2018). We compared the 3′ UTR sequences of the HNRNPA2B1-target RNAs to 3′ UTR sequences from thousands of RNAs whose localization was not sensitive to the loss of HNRNPA2B1 (**Figure 1E**). We found that the 3′ UTRs of HNRNPA2B1-target RNAs were highly enriched for AG-rich sixmers, suggesting that HNRNPA2B1 directly binds these RNAs to regulate their localization (**Figure 3A, 3B**). This effect was muted when comparing coding sequences from the two sets of RNAs (**Figure S2A**) and completely lost when comparing 5′ UTR sequences (**Figure S2B**). Given that HNRNPA2B1 predominantly binds to 3′ UTR sequences (Huelga et al. 2012; Martinez et al. 2016) and many localization-regulatory sequences are found in 3′ UTRs (Engel et al. 2020), this further supports the hypothesis that HNRNPA2B1 is binding the target RNAs to regulate their localization.

**Figure 3.**
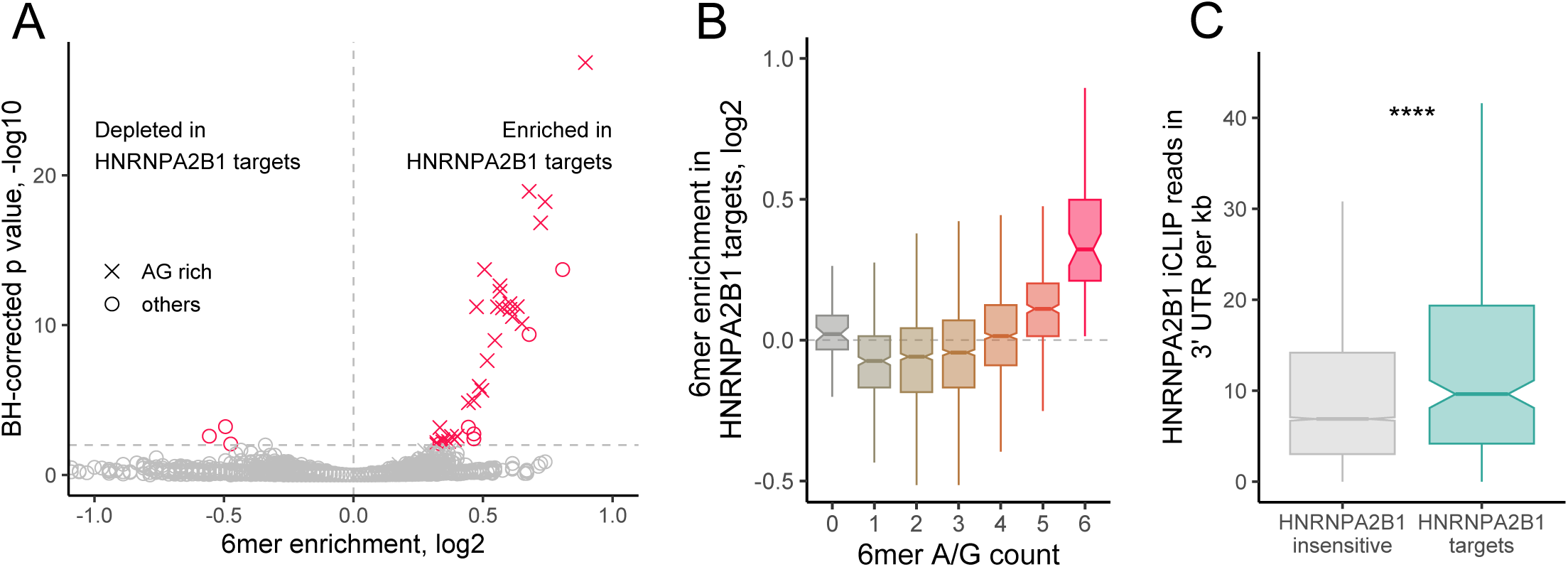
HNRNPA2B1 localization targets are enriched for HNRNPA2B1 motifs and CLIP sites in their 3′ UTRs. (A) Enrichments for all 6mers in the 3′ UTRs of HNRNPA2B1 targets RNAs compared to the 3′ UTRs of RNAs of HNRNPA2B1-insensitive RNAs. 6mers that are completely comprised of A and G are represented with X. All others are represented with circles. (B) Distributions of 6mer enrichments in the 3′ UTRs of HNRNPA2B1 targets RNAs compared to the 3′ UTRs of RNAs of HNRNPA2B1-insensitive RNAs. 6mers are binned according to the number of A/G they contain. (C) HNRNPA2B1 CLIP read densities in the 3′ UTRs of HNRNPA2B1-insensitive (gray) and HNRNPA2B1-target (teal) RNAs. P-values were calculated using a Wilcoxon ranksum test. NS (not significant) represents p > 0.05, * p < 0.05, ** p < 0.01, *** p < 0.001, and **** represents p < 0.0001.

To further ask if HNRNPA2B1-target RNAs were directly bound by HNRNPA2B1 in cells, we analyzed available CLIP data for the RBP derived from mouse spinal cord samples (Martinez et al. 2016). We found that the 3′ UTRs of the HNRNPA2B1-target genes had significantly higher read densities in the CLIP data than nontargets (**Figure 3C**). They were also approximately 2.5 times more likely to contain CLIP read peaks than nontargets (**Figure S2C**), further suggesting that the HNRNPA2B1-target RNAs were directly bound by HNRNPA2B1 in cells. This effect was again muted when comparing coding sequences and completely lost when comparing 5′ UTR sequences (**Figure S2D, S2E**).

To test if the localization regulation of the target RNAs may be conserved in human cells, we compared the human orthologs of the HNRNPA2B1 targets to the human orthologs of the nontargets. The 3′ UTRs of the human ortholog targets were also highly enriched in AG-rich sequences (**Figure S2F, S2G**). We then analyzed HNRNPA2B1 CLIP data from human cells (Nguyen et al. 2018). We found that the 3′ UTRs of the human orthologs of the HNRNPA2B1 targets were significantly more likely to contain CLIP peaks than the orthologs of the nontargets, suggesting that these targets by also be subject to regulation by HNRNPA2B1 in human cells (**Figure S2H**).

### Arginine to alanine mutations in the RGG domain of HNRNPA2B1 result in its increased cytoplasmic distribution

To further probe connections between HNRNPA2B1 levels and the localization of HNRNPA2B1-target RNAs, we devised a strategy whereby we modulated the levels of HNRNPA2B1 in the cytoplasm. Since RNA transport to neurites occurs in the cytoplasm, we reasoned that if HNRNPA2B1 was truly regulating the localization of these RNAs, increasing its cytoplasmic levels would cause its localization-regulatory effects to become stronger.

HNRNPA2B1 contains an RGG domain (**Figure 4A**). These domains are rich in arginine and glycine and are found in many RBPs where they often regulate nucleocytoplasmic trafficking of the RBP (Thandapani et al. 2013; Chowdhury and Jin 2023). This trafficking is often regulated by methylation of arginine residues within the RGG domain of the RBP (Dormann et al. 2012; Scaramuzzino et al. 2013; De Leeuw et al. 2007).

**Figure 4.**
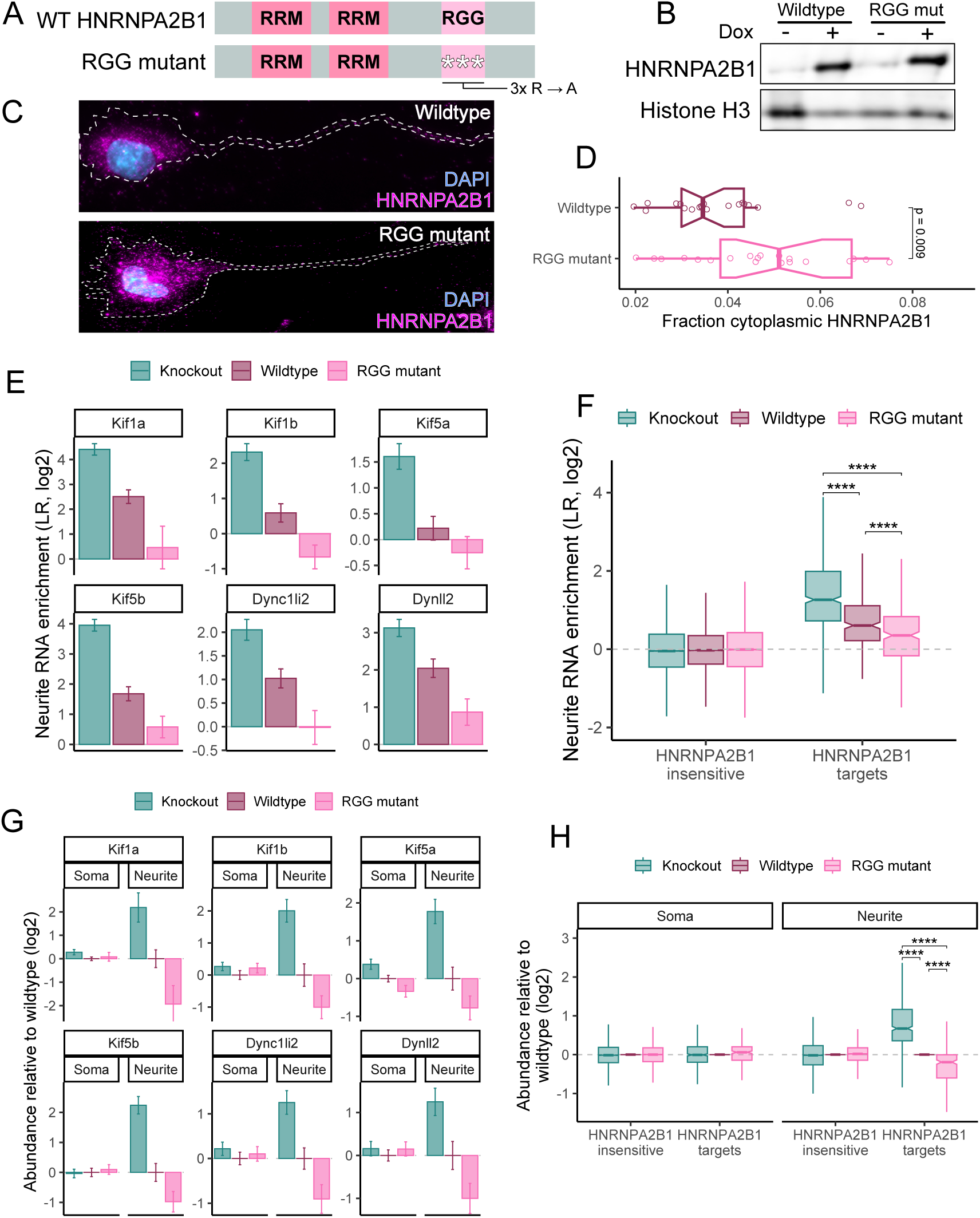
Mutations in the RGG domain of HNRNPA2B1 make it more cytoplasmic and a more potent regulator of neurite RNA localization. (A) Domain organization of HNRNPA2B1 rescue constructs. RNA recognition motifs are designated with RRM. (B) Immunoblot of wildtype and RGG mutant HNRNPA2B1 rescue transgenes. (C) Immunofluorescence of wildtype and RGG mutant HNRNPA2B1 rescue transgenes. (D) Quantification of the fraction of HNRNPA2B1 localized to the cytoplasm with wildtype and RGG mutant rescue transgenes. (E) Neurite enrichment levels of select motor protein RNAs in HNRNPA2B1 knockout, wildtype rescue, and RGG mutant rescue cells. (F) Neurite enrichment levels for HNRNPA2B1-insensitive and HNRNPA2B1-target RNAs (as defined in figure 1E) in HNRNPA2B1 knockout, wildtype rescue, and RGG mutant rescue cells. (G) RNA expression levels for select motor proteins in the soma and neurites of HNRNPA2B1 knockout, wildtype rescue, and RGG mutant rescue cells. Values are shown relative to expression levels in wildtype rescue cells. (H) Compartment-specific RNA expression levels for HNRNPA2B1-insensitive and HNRNPA2B1-target RNAs (as defined in figure 1E) in HNRNPA2B1 knockout, wildtype rescue, and RGG mutant rescue cells. P-values were calculated using a Wilcoxon ranksum test. NS (not significant) represents p > 0.05, * p < 0.05, ** p < 0.01, *** p < 0.001, and **** represents p < 0.0001.

Like several other RGG-containing proteins, the localization of HNRNPA2B1 is dependent upon the methylation status of arginine residues in its RGG domain (Nichols et al. 2000). Loss of either methylation activity of the RGG domain itself causes HNRNPA2B1 to shift from a primarily nuclear distribution to a primarily cytoplasmic distribution (Nichols et al. 2000). We therefore hypothesized that mutation of arginine residues within the RGG domain of HNRNPA2B1 would result in an increase in its cytoplasmic localization.

To test this, we rescued our HNRNPA2B1 knockout cells with either wildtype HNRNPA2B1 or a mutant in which three arginine residues in the RGG domain were mutated to alanine (**Figure 4A**). The arginine mutations did not affect the total expression levels of the rescues (**Figure 4B**). However, they did result in significantly increased levels of HNRNPA2B1 in the cytoplasm (**Figure 4C, 4D**), consistent with earlier reports (Nichols et al. 2000). With this system, we now had the means to titrate the cytoplasmic levels of HNRNPA2B1 from low expression levels of the rescues (**Figure 4B**). However, they did result in significantly increased levels of HNRNPA2B1 in the cytoplasm (**Figure 4C, 4D**), consistent with earlier reports (Nichols et al. 2000). With this system, we now had the means to titrate the cytoplasmic levels of HNRNPA2B1 from low (knockout) to moderate (wildtype rescue) to high (RGG mutant rescue).

### Increasing the amount of HNRNPA2B1 in the cytoplasm increases its RNA localization regulatory ability

We hypothesized that increasing the cytoplasmic level of HNRNPA2B1 would increase its ability to regulate RNA abundance in neurites. To test this, we profiled soma and neurite RNA levels using subcellular fractionation and high-throughput RNA sequencing in HNRNPA2B1 knockout, wildtype rescue, and RGG mutant rescue CAD cells. When comparing knockout and wildtype rescue samples, we again found that many RNAs encoding motor proteins were significantly more neurite enriched in knockout cells compared to wildtype rescue cells. In RGG mutant rescue cells, the neurite enrichment of these RNAs was decreased even further beyond that seen in wildtype rescue cells, consistent with the RGG mutant rescue more potently regulating RNA localization than the wildtype rescue (**Figure 4E, Table S3**).

This effect was not limited to these specific examples. We separated RNAs into HNRNPA2B1-target RNAs and HNRNPA2B1-insenstive RNAs as defined by our first subcellular RNA sequencing experiment (**Figure 1E**). While HNRNPA2B1-insensitive RNAs were again insensitive to loss and/or HNRNPA2B1 rescue, rescue of the knockout cells with the RGG mutant reduced the neurite enrichment of of HNRNPA2B1-target RNAs beyond what was observed with rescue with wildtype HNRNPA2B1 (**Figure 4F**). These results suggest that increasing cytoplasmic levels of HNRNPA2B1 allow it to more potently regulate RNA localization to neurites.

We then looked at soma and neurite RNA levels separately for knockout, wildtype rescue, and RGG mutant rescue cells. As before, RNA levels for several motor proteins were generally unchanged in the soma between knockout, wildtype rescue and RGG mutant rescue samples. However, RNA levels in neurites were significantly affected. Relative to wildtype rescue cells, neurite RNA levels were increased in knockout cells and decreased in RGG mutant cells (**Figure 4G**). This effect was again seen throughout the group of HNRNPA2B1-target RNAs (**Figure 4H**). In total, these results indicate a dose-dependent connection between the cytoplasmic levels of HNRNPA2B1 and the neurite-specific levels of HNRNPA2B1-target RNAs.

### HNRNPA2B1 loss inhibits motor protein activity specifically in neurites

Given that the neurite-specific levels of many mRNAs encoding motor proteins were aberrantly increased in HNRNPA2B1 knockout cells, we wondered if neurite-specific motor protein function was also affected. To assess this, we monitored the transport of lysosomes in live cells using a lysosome-specific fluorescent dye (Barral et al. 2022). While in wildtype rescue cells lysosomes in neurites were highly motile, in HNRNPA2B1 knockout cells neurite lysosomes were markedly more static. RGG mutant HNRNPA2B1, which was competent for regulating RNA localization, was able to rescue the motor protein activity defect (**Figure 5A, Videos S1-S3**).

**Figure 5.**
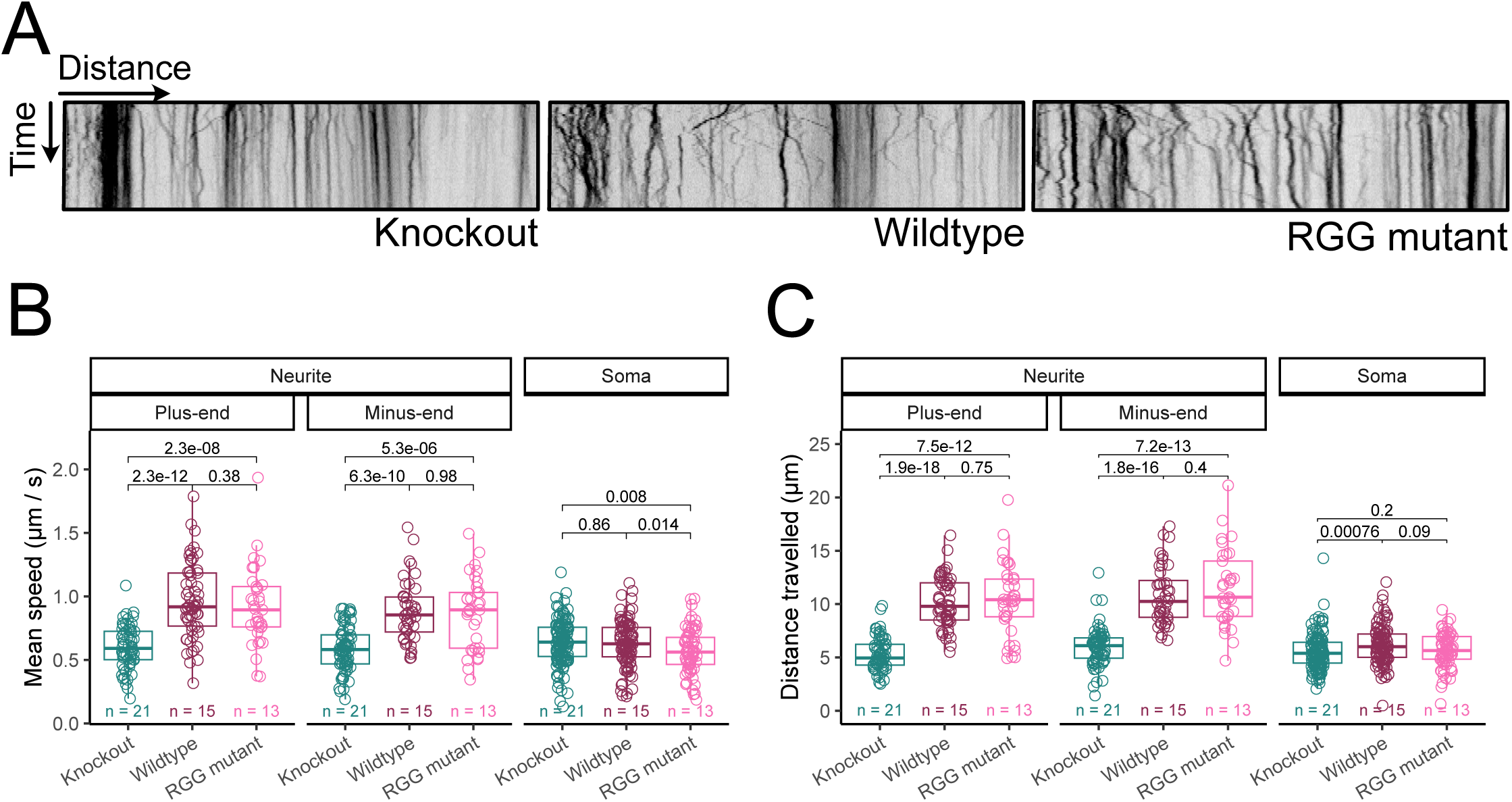
Motor protein activity is impaired in the neurites of HNRNPA2B1 knockout cells and accelerated in cells expressing RGG mutant HNRNPA2B1. (A) Kymographs of lysosome movement in the neurites of HNRNPA2B1 knockout (left), wildtype rescue (middle) and RGG mutant rescue (right) cells. (B) Quantification of lysosome trafficking speed. The number of cells assayed in each condition is specified. (C) Quantification of lysosome trafficking distance. The number of cells assayed in each condition is specified. P-values were calculated using a Wilcoxon ranksum test.

We calculated velocities for individual lysosomes in neurites traveling towards the plus-end (towards the tip of the neurite) and minus-end (towards the soma) directions. We also calculated velocities for lysosomes in somas but were unable to assign directionality to them due to the less defined cytoskeletal organization in cell bodies. We found that in wildtype rescue cells, lysosome traveled approximately 0.5-1.0 µm / sec, consistent with previous reports (Bandyopadhyay et al. 2014).

In HNRNPA2B1 knockout cells, both plus- and minus-end directed velocities were significantly reduced in the neurite while their velocities in the soma were much less affected (**Figure 5B**). Similarly, relative to wildtype rescue cells, the distances traveled by individual lysosomes in neurites were significantly lower in knockout cells. This effect was again generally neurite-specific as the magnitude of differences was greatly reduced in the soma (**Figure 5C**). The abundance of mRNAs encoding motor proteins and the activities of those proteins are therefore misregulated in the neurites of HNRNPA2B1 knockout cells, although a mechanistic connection between those observations remains unknown.

### The stability of HNRNPA2B1 RNA localization targets is regulated by the deadenylase CAF1

Although these results indicate that the neurite abundance of hundreds of mRNAs are regulated by HNRNPA2B1, they don’t explain how that is happening. The phenomenon of an RBP negatively regulating neurite RNA abundance has been reported in the literature previously only once, for the RBP Pum2 (Martínez et al. 2019). Interestingly, both Pum2 and HNRNPA2B1 have been found to negatively regulate RNA stability (Goldstrohm et al. 2018; Geissler et al. 2016). Loss of these proteins would therefore be expected to increase the stability of their targets. Recent work has found that increasing an RNA’s stability is associated with an increase of its abundance in neurites (Loedige et al. 2023). We therefore hypothesized that an increase in the stability of HNRNPA2B1-target RNAs following the loss of HNRNPA2B1 leads to their increased neurite abundance.

To test this, we began with the reported observation that HNRNPA2B1 facilitates RNA decay by recruiting the CCR4-NOT deadenylase complex (Geissler et al. 2016). One of the enzymatic subunits of this deadenylase is CAF1. A recent study quantified changes in the stability and neurite enrichment of RNAs in mouse cortical neurons following the inactivation of CAF1 through the expression of a dominant negative version of the protein (Loedige et al. 2023).

If our hypothesis that the RNAs whose localization is regulated by HNRNPA2B1 are also destabilized by HNRNPA2B1 is correct, then our HNRNPA2B1 localization targets should be preferentially stabilized following the loss of CCR4-NOT deadenylase activity. We found that, indeed, compared to all RNAs, HNRNPA2B1 localization targets were preferentially stabilized following the loss of CCR4-NOT deadenylase activity (**Figure 6A**).

**Figure 6.**
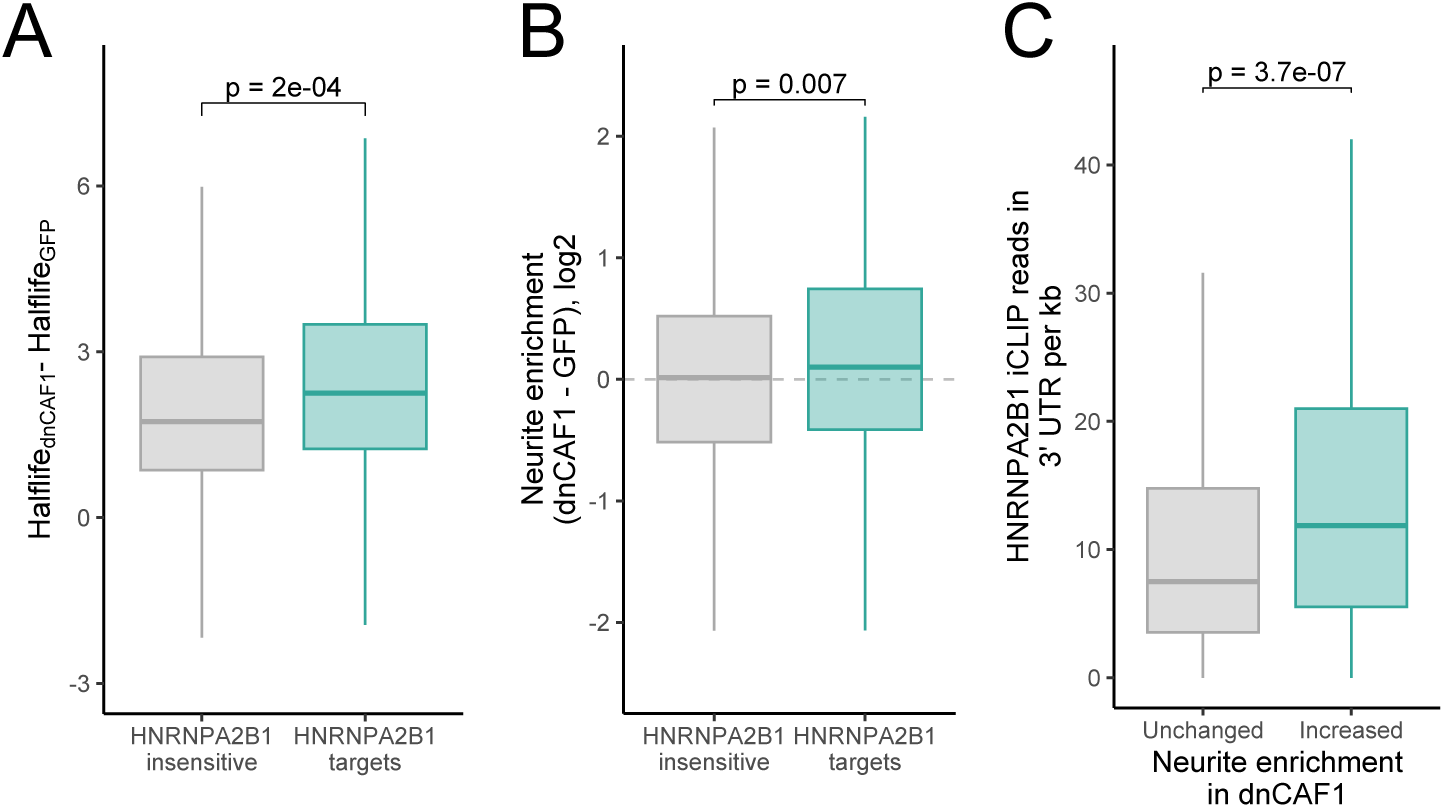
The stability and localization of HNRNPA2B1 localization targets is sensitive to loss of deadenylase activity. (A) Changes in RNA halflife between cells expressing dominant negative CAF1 and those expressing GFP as a control for both HNRNPA2B1-insensitive RNAs and HNRNPA2B1-target RNAs (as defined in figure 1F). (B) Changes in neurite enrichment between cells expressing dominant negative CAF1 and those expressing GFP as a control for both HNRNPA2B1-insensitive RNAs and HNRNPA2B1-target RNAs. (C) Density of HNRNPA2B1 CLIP-seq reads in the 3′ UTRs of RNAs whose neurite-enrichment was unchanged (gray) and increased (teal) in response to the inhibition of deadenylation through the expression of dominant negative CAF1.

### The neurite-enrichment of HNRNPA2B1-target RNAs is sensitive to the loss of CCR4-NOT deadenylase activity

We hypothesized that the loss of HNRNPA2B1-mediated destabilization via CCR4-NOT recruitment of its target RNAs leads to their increased abundance in neurites. If this is correct, then the RNAs that become more neurite-enriched following HNRNPA2B1 loss should also become more neurite-enriched following the loss of CCR4-NOT deadenylase activity. This is indeed what we observed as, compared to HNRNPA2B1-insensitive RNAs, our HNRNPA2B1 localization-target RNAs became significantly more neurite-enriched following the loss of CCR4-NOT deadenylase activity (**Figure 6B**).

Finally, if HNRNPA2B1’s ability to negatively regulate RNA stability via CCR4-NOT recruitment contributes to its ability to regulate neurite enrichment, we would expect that RNAs whose neurite-enrichment increases upon CCR4-NOT activity loss to be enriched for HNRNPA2B1 binding. This is what we observed as RNAs that became more neurite-enriched following CAF1 inactivation were significantly enriched for HNRNPA2B1 CLIP-seq reads in their 3′ UTRs compared to CAF1-insensitive RNAs (**Figure 6C**). In sum, although these data do not prove that HNRNPA2B1 regulates neurite RNA localization via regulating RNA stability, they are consistent with the idea.

## DISCUSSION

Although hundreds of RNAs are known to be enriched in neuronal projections, the molecular mechanisms that are regulating this localization are unknown for the vast majority. In this study we have identified RNAs whose proper localization depends on the RBP HNRNPA2B1. Unexpectedly, we found that HNRNPA2B1 mainly acts to keep RNAs out of neurites as its loss leads to significantly increased neurite abundances for many transcripts. These HNRNPA2B1-sensitive RNAs were enriched for HNRNPA2B1 sequence motifs and CLIP-derived binding sites and often encoded motor proteins. In HNRNPA2B1 knockout cells, the activity of motor proteins was impaired specifically in neurites. Interestingly, the ability of HNRNPA2B1 to regulate neurite RNA accumulation and motor protein activity was enhanced by increasing its cytoplasmic abundance through the incorporation of specific point mutations. These data allow us to construct a model in which HNRNPA2B1 binds RNAs in their 3′ UTRs to prevent their accumulation in neurites, perhaps through regulating their stability. Upon the loss of HNRNPA2B1, these transcripts aberrantly accumulate in neurites, somehow leading to the impairment of motor protein activity specifically in neurites (**Figure 7**).

**Figure 7.**
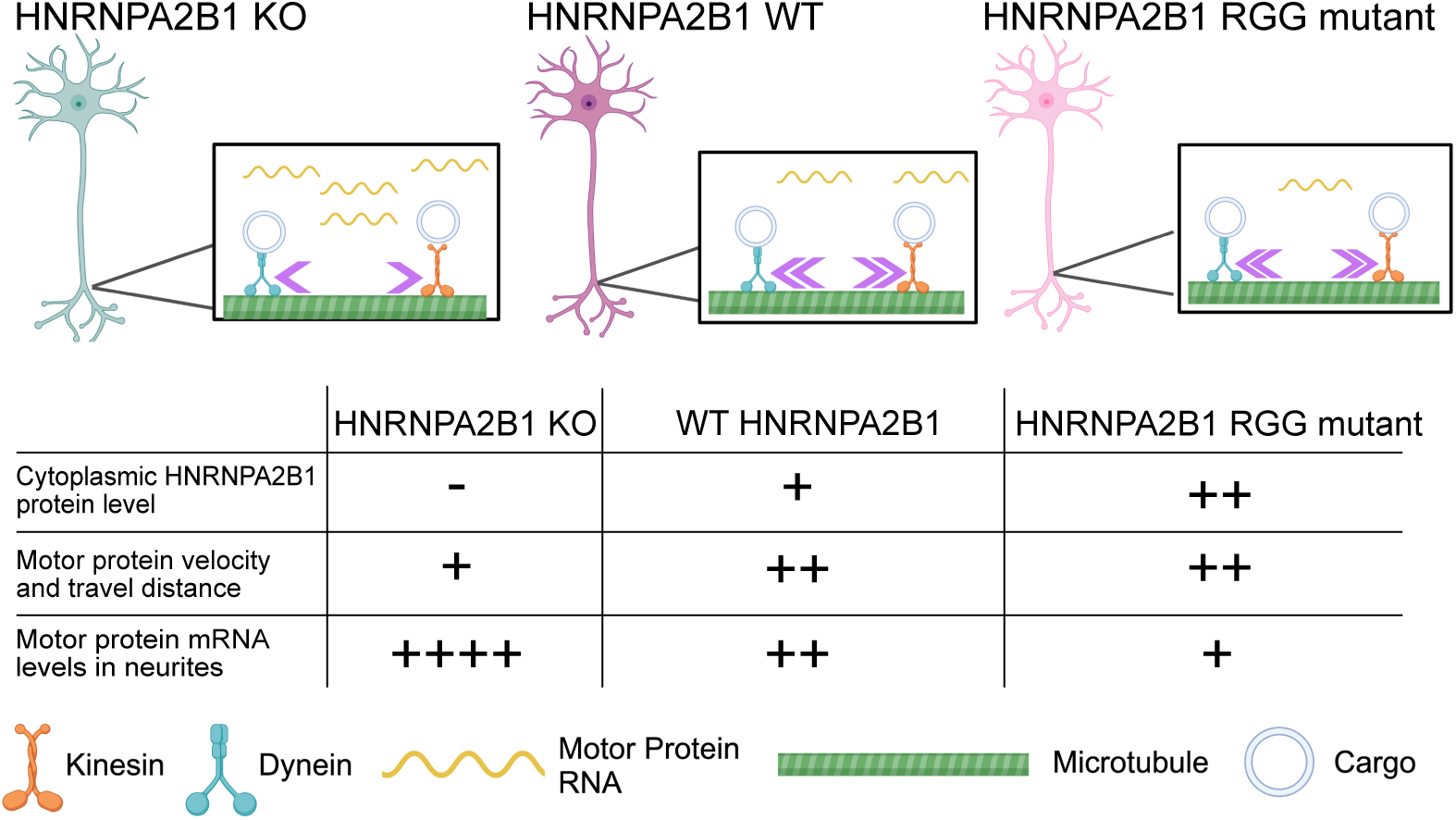
Model. In wildtype cells (middle), RNAs encoding motor proteins are present at moderate levels in neurites, and motor protein activity is normal. In HNRNPA2B1 knockout cells (left), motor protein RNA levels are increased specifically in neurites, and motor protein activity in neurites is decreased. In cells expressing RGG mutant HNRNPA2B1, motor protein RNA levels in neurites are decreased below their levels in wildtype cells, and motor protein activity in neurites is restored. The presence and/or direction of causality between motor protein RNA levels in neurites and motor protein activity in neurites remains unknown.

It is important to note that we only have correlative, rather than direct, evidence that an increase in motor protein mRNAs in neurites is associated with a loss in motor protein activity at that location. Future studies will be required to thoroughly investigate any causality. However, it has been demonstrated that the location of motor protein mRNA translation can have effects on trafficking-related phenotypes (Norris and Mendell 2023). It is therefore at least plausible that these observations are directly mechanistically connected.

Reduced levels of functional HNRNPA2B1 and point mutations within the protein have been linked to multiple neurodegenerative diseases, including Alzheimer’s disease and amyotrophic lateral sclerosis (ALS) (Martinez et al. 2016; Kim et al. 2013; Berson et al. 2012). One pathogenic mutation, D290V, results in HNRNPA2B1 being localized to cytoplasmic inclusions (Martinez et al. 2016). Given that axon degeneration is a key phenotype in many neurodegenerative disorders, it is tempting to conceptually connect HNRNPA2B1 levels and/or localization, the loss of motor protein activity in axons, and neurodegenerative phenotypes. However, no evidence for this direct connection currently exists.

Although earlier studies identified HNRNPA2B1 as promoting RNA transport to cellular projections (Ainger et al. 1997; Shan et al. 2000), our results indicate a mainly repressive role in RNA transport. This could be because the earlier studies focused only on one particular *cis-*element that was both bound by HNRNPA2B1 and promoted transport to projections. Alternatively, and not mutually exclusively, it may be because those studies focused on oligodendrocyte cell types rather than neuronal cell types.

If HNRNPA2B1 regulates neuronal RNA localization through regulating RNA stability, this opens up the possibility that many RBPs may be similarly affecting RNA trafficking in neurons. Dozens of proteins regulate RNA stability (Houseley and Tollervey 2009), and so the collections of RBPs that, directly or indirectly, regulate RNA localization may be similarly large. Given that the RBP-directed regulatory landscape of RNA localization is perhaps not as understood as that of other RNA-based regulatory processes, this means that much work remains to be done in how RNAs end up at where they are supposed to be.

## ACKNOWLEDGEMENTS

We thank the Taliaferro and Moore labs for helpful and productive discussions and Lisa Wood for instruction and guidance regarding particle tracking in live cells. This work was funded by NIH grants R35GM133385 (J.M.T.), R35GM136253 (J.K.M.), and R01NS122911 (J.M.T. and H.A.R.).

## METHODS

### CAD cell maintenance and differentiation

Prior to integration of transgenes, CAD cell lines were maintained in 1:1 DMEM/F12 mix (Thermo 11320033), 10% Equafetal (Atlas Biologicals EF-0500-A), 5.0 µg/mL blasticidin (A.G. Scientific B-1247-SOL), and 0.01% penicillin-streptomycin (Thermo 15140122). After integration of transgenes, blasticidin was omitted and 5.0 µg/ mL puromycin (Cayman Chemical Company 13884) was added. 2 µg/mL doxycycline was added for 48 hrs to induce expression of rescue constructs. To differentiate CAD cells, full growth media was replaced with serum free media to induce neurite outgrowth.

### Generation of an Hnrnpa2b1 KO cell line with a single loxP cassette

CAD cell lines were authenticated with STR profiling and found to be mycoplasma negative. The guide RNA (gRNA) sequences AGAAACUACUAUGAGCAAUG and AUUGCAGGGAACUAUGGUCC for *Hnrnpa2b1* were obtained from Synthego and dissolved in 1x TE buffer for a final concentration of 100 µM. Purified Cas9 protein (Synthego) was added to the two gRNAs at a 3:1 molar ratio of gRNA:Cas9 at room temperature for 15 minutes. Then, these RNP complexes were co-electroporated with a GFP encoding plasmid into CAD cells using the Neon transfection system (ThermoFisher) with the following settings - 1400 V, 1 pulse for 30 ms. 48 hours after transfection, cells were sorted to single cells with a flow cytometer and allowed to grow for two weeks. Clones were then screened for HNRNPA2B1 expression with a polyclonal Hnrnpa2b1 antibody. Lesions at the *Hnrnpa2b1* locus were identified by PCR from genomic DNA from single cell clones and screened for *Hnrnpa2b1* deletion.

### Expression of HNRNPA2B1 rescue constructs in CAD cells

CAD *Hnrnpa2b1* KO cells were plated in a 6 well plate at approximately 6 x 10^5^ per well in DMEM:F12 media supplemented with 10% Equafetal for 48 hours before transfection. Cells were co-transfected with a plasmid derivative containing Hnrnpa2b1 rescue cDNs with 1% wt/wt of a Cre encoding plasmid (pCAGGS-Cre). For the transfection of one well of a 6 well plate, 0.5 µg of HNRNPA2B1-encoding plasmid was mixed with 20 ng of Cre-encoding plasmid, 8 µL of Lipofectamine LTX, 4 µL PLUS reagent and 200 µL Optimem based on the manufacturer’s protocol. Cells were incubated overnight with the transfection reagents before the media was replaced and cells were incubated for an additional 24 hours. 5.0 µg/mL puromycin was added until cells in untransfected wells were depleted. Remaining cells with stably integrated rescue constructs were expanded in full growth media. Lysates were then collected to assess rescue construct expression by western blot and immunofluorescence.

### Immunoblotting for Hnrnpa2b1

*Hnrnpa2b1* KO CAD cells with integrated rescue constructs were plated in 6 well plates at 1.08 x 10^6^ cells per well in duplicate in full growth media or full growth media supplemented with 2.0µg/mL doxycycline. Lysates were collected in 50 µL of ice cold RIPA buffer, incubated on ice with periodic vortexing for five minutes before centrifugation at 21000 rcf for 5 minutes. The supernatant was removed and 15µL of the lysate was added to sample buffer (Invitrogen NP008) and DTT at a final concentration of 3 mM and boiled at 95℃ for five minutes. Remaining lysate was stored at −80°C. Denatured lysates were separated by PAGE on 4%-12% Bis-Tris gradient along with Spectra Protein Ladder (Thermo Fisher 26634) in MOPS SDS Running Buffer (ThermoFisher NP0001) at 150V for 90 minutes. The gel was transferred to a PVDF membrane using an iBlot2 dry transfer machine (Thermo Fisher IB21001). Total protein was assessed with Ponceau staining and destained before blots were blocked with agitation in 5% milk powder in PBST for 45 minutes at room temperature. Blots were subsequently washed 3 times for 5 minutes each in PBST at room temperature before incubation in 5µg/mL of HNRNPA2B1 polyclonal primary antibody (abcam 31645) in PBST overnight at 4°C with agitation. Blots were washed three times for 5 minutes each with agitation and incubated with anti-rabbit IgG HRP-conjugated secondary antibody (Proteintech SA00001-2) for 30 minutes at room temperature. Blots were washed 3 times for 5 minutes in PBST and visualized with the WesternBright Sirius HRP kit on a Sapphire imager (Azure Biosystems) set to collect chemiluminescence signal.

### Immunofluorescence for HNRNPA2B1

CAD *Hnrnpa2b1* KO cells with doxycycline inducible HNRNPA2 rescue constructs were plated on PDL coated glass coverslips (neuVitro) that fit within 12 well plates at 4.0 x 10^4^ cells per well in full growth media or full growth media with 2.0 µg/mL doxycycline for 3 hours to allow cells to attach to the coverslip. Media was removed and replaced with serum free media or serum free media with 2.0µg/mL doxycycline to induce neurite differentiation for 48 hours. Cells were rinsed twice with PBS for 5 minutes before fixation with 3.7% formaldehyde and permeabilization with 0.1% Triton-X in PBS for 12 minutes at room temperature. Cells were blocked with 1% BSA, 0.1% Triton-X and 75 mM NaCl in PBS for 30 minutes with agitation. Cells were rinsed twice with PBS before primary HNRNPA2B1 antibody (abcam 31645) was added at a 1:1000 dilution in PBS containing 1% BSA and 0.1% Triton-X overnight at 4°C. Cells were rinsed 3 times with PBS for 5 minutes before adding fluorescent secondary antibody (Cell Signaling Technology 4413S) diluted 1:2000 in PBS containing 1% BSA and 0.1% Triton-X at 37°C for 25 minutes. After this solution was removed, cells were incubated with 100 ng/mL DAPI in PBS containing 1% BSA and 0.1% Triton-X at 37°C for 5 minutes. Cells were rinsed 3 times with PBS before mounting onto slides with Fluoromount G and sealed with nail polish.

### Fractionation and sequencing of subcellular transcriptomes

Fractionation of neuronal cells were performed as previously described (Arora et al. 2021). CAD cells were plated on porous transwell membranes with a pore size of 1.0 micron (Corning 353102) at a density of 1 million cells per membrane which fit in one well of a 6 well plate. Each 6 well plate comprised one biological replicate. After cells were attached, cells were differentiated in serum free media to induce neurite formation for 48 hours.

To fractionate CAD cells into soma and neurite fractions, membranes were first washed with PBS before 1 mL of PBS was added on top of the membrane. The top of the membrane, containing the soma, was repeatedly scraped with a cell scraper to remove the soma and placed into an ice cold 15 mL falcon tube. Following scraping, membranes were removed from its plastic housing using a razor blade and placed in 550 µL of RNA lysis buffer (Zymo R1050) at room temperature for 15 minutes to lyse the neurites. Soma samples were spun down and resuspended in 300µL PBS. 100 µL of this soma sample was carried forward for RNA isolation with the Zymo QuickRNA MicroPrep Kit (Zymo R1050). RNA was isolated from the 500 µL of neurite lysate in parallel with the same kit. The expected neurite RNA yield from six membranes was 100-300 ng.

PolyA selected, stranded libraries were synthesized using 100 ng total RNA with the KAPA mRNA Hyperprep Kit (KAPA/Roche). Libraries were sequenced with paired end 150 bp sequencing on an Illumina sequencer. Approximately 40 million read pairs were obtained per sample. Each condition (HNRNPA2B1 KO, HNRNPA2B1 rescue, soma and neurite) was collected and sequenced in triplicate.

### Taqman qPCR of subcellular compartment RNA samples

cDNA was synthesized from 50 ng of RNA from each compartment using the LunaScript RT supermix kit (NEB #E3010) using the manufacturer’s protocol. The cDNA was diluted to 20 µL with RNAse free water. 2 µL of diluted cDNA was used as the template to assess the quality of the soma/neurite fractionation with qPCR using Taqman mastermix (IDT 1055772) to quantify endogenous *Tsc1* and *Net1* RNAs. *Net1* RNA is reproducibly highly neurite-enriched and therefore if a soma/neurite fractionation is successful, should be more relatively abundant in neurites than the nonlocalized transcript *Tsc1*. The relative expression of these genes were quantified within the same sample to calculate log_2_ normalized localization ratios. All qPCRs were performed in triplicate averaged to produce a single biological replicate.

### Quantification of RNA localization from CAD subcellular RNA-sequencing data

Transcript abundances were calculated using salmon v1.5.2 (Patro et al. 2017) using the Gencode M17 genome annotation for *Mus musculus*. Transcript abundances were collapsed to gene abundances with txImport (Soneson et al. 2015). Localization ratios were calculated for each gene as the log_2_ of the ratio of normalized counts in neurite/soma for a gene by DESeq2 using the salmon/txImport calculated counts (Love et al. 2014).

### smFISH visualization of RNA localization

CAD HNRNPA2B1 knockout and rescue cells were plated on PDL coated glass coverslips (neuVitro) that fit within 12 well plates at 4.0 x 10^4^ cells per well in full growth media or full growth media with 2.0 µg/mL doxycycline for 3 hours to allow cells to attach to the coverslip. Media was removed and replaced with serum free media or serum free media with 2.0 µg/mL doxycycline (rescue) to induce neurite differentiation for 48 hours. The media was aspirated, and cells were washed once with 1× PBS. Cells were fixed in 3.7% formaldehyde (Fisher Scientific) in PBS for 10 min at room temperature and then washed twice with 1× PBS. Cells were permeabilized with cold 70% ethanol (VWR) for 2 hours at 4°C. The cells were washed with freshly prepared wash buffer (2X SSC and 10% formamide in water) at room temperature for 15 minutes.

In the meantime, the smiFISH probes for each gene were hybridized to the fluorescent Y Flap using the protocol from (Tsanov et al. 2016). *Kif5b* and *Kif1a* probe sets included 48 probes, and *DynII2* contained 36 probes. After hybridization, the probe/flap hybridization product was spun in a benchtop centrifuge for 60 seconds. Per coverslip, 2 µL (0.833 µM) of smFISH probe was added to 100 µL of smFISH hybridization buffer (Biosearch Technologies) for each condition and gene. A hybridization chamber was prepared using an empty tip container, wrapped in tinfoil with parafilm and wet paper towels inside the box to retain moisture. 100 µL of the probe-containing hybridization solution was added to the parafilm. The coverslip was then placed on top of this droplet of hybridization buffer with the cell side down. The hybridization chamber with the coverslips was incubated at 37°C overnight (15-18 hours). The coverslips were transferred to a fresh 12-well plate with the cell side up and incubated twice with freshly prepared wash buffer for 45 minutes at 37°C. The second wash included a membrane dye (CellBrite® Fix 488). Then slides were incubated with wash buffer including DAPI (100 ng/mL) at 37°C for 30 minutes. Slides were washed twice with PBS for 5 minutes at room temperature. Coverslips were then mounted onto slides with Fluoromount G and sealed with nail polish.

Slides were imaged at 63x with consistent laser intensity and exposure times across samples. DAPI was imaged with an exposure of 100 ms. Membrane dye was used to find cells and was imaged in the FITC channel with an exposure of 100ms. FISH probes were visualized in the TRITC channel with an exposure of 1500ms. Z stacks were collected of approximately 24 images, 0.4 µm apart.

### Quantification of smFISH images

FISH-quant was used to quantify neuronal projection enrichment of smFISH spots as previously described (Tsanov et al. 2016; Arora et al. 2022b). Briefly, outlines were drawn in the FITC channel visualizing the cell membrane. Two outlines were drawn per cell: soma and neuronal projection. Prior to quantification, identified smFISH spots were thresholded for intensity, sphericity, amplitude, and position. Transcript enrichment was quantified by the total number of spots in the projection of cells over a total number of spots in the soma.

### Identification of enriched RNA motifs

Transcript region (5′ UTR, CDS, 3′ UTR) sequences were derived for all transcripts using Gencode vM17 annotations. For each gene, the longest transcript region was used. Genes were binned according to their neurite-enrichment sensitivity to HNRNPA2B1 loss. Enriched kmers were identified using custom Python scripts.

### Comparison to HNRNPA2B1 CLIP-seq data

HNRNPA2B1 CLIP data from mouse spinal cord samples was downloaded (Martinez et al. 2016). Adapters were removed using the following cutadapt call: cutadapt --match-read-wildcards --times 2 -e 0 -O 5 --quality-cutoff 6 -m 18 -b TCGTATGCCGTCTTCTGCTTG -b ATCTCGTATGCCGTCTTCTGCTTG -b CGACAGGTTCAGAGTTCTACAGTCCGACGATC -b TGGAATTCTCGGGTGCCAAGG -b AAAAAAAAAAAAAAAAAAAAAAAAAAAAAAAAAAAAAAAAAAAAAAAAAA -b TTTTTTTTTTTTTTTTTTTTTTTTTTTTTTTTTTTTTTTTTTTTTTTTTT (Martin 2011). Reads were then aligned to the mouse genome using STAR (Dobin et al. 2013). Duplicate reads were masked using samtools (Li et al. 2009), and reads were intersected with transcripts using bedtools (Quinlan and Hall 2010).

### Quantifying nuclear and cytoplasmic HNRNPA2B1 protein abundance in CAD cells

The nuclear/cytoplasmic distribution of HNRNPA2B1 was quantified using the masking approach as previously reported (Kelley and Paschal 2019). The nuclear marker DAPI was used to create a nuclear mask for individual cells. The cytoplasm was defined by selecting the cell area using the segmented line tool in Fiji (Schindelin et al. 2012). The nuclear-cytoplasmic ratio of HNRNPA2B1 was calculated by dividing the mean nuclear intensity by the mean cytoplasmic intensity.

### Visualizing lysosome dynamics in live CAD cells

CAD HNRNPA2B1 KO cells with doxycycline inducible HNRNPA2 rescue constructs were plated on glass petri dishes at approximately 0.325 x 10^6^ cells per dish in full growth medium or full growth media supplemented with 2.0 µg/mL doxycycline for 3 hours to enable cell attachment. Media was removed and replaced with serum free medium and supplemented with 2.0 µg/mL doxycycline for 48 hours to induce neurite differentiation and rescue construct expression. Media was removed and washed with PBS before incubation in full growth media with 1.0µg/mL Lysoview 540 (Biotium) for 25 minutes at 37°C. Media was replaced before imaging with a Nikon CSU-W1 SoRa Spinning Disk Confocal Microscope at 20X with consistent laser power and exposure times across samples. Images were collected on a Nikon Ti2-E inverted microscope equipped with 1.45 NA 100x CFI Plan Apo objective (Nikon Inc; Melville, NY), Nikon motorized stage, Prior NanoScan SP 600 mm nano-positioning piezo sample scanner (Prior Scientific; Rockland, MA), CSU-W1 T1 Super-Resolution spinning disk confocal, SoRa disk with 1x/2.8x/4x mag changers (3i; Denver, CO), 488 nm, 560 nm, and 647 nm laser, and a prime 95B back illuminated sCMOS camera (Teledyne Photometrics; Tuscon, AZ). Temperature was maintained at 37°C and 5% CO_2_ using a chamber surrounding the microscope stage. The 540 channel was visualized with 80% laser power for 300 ms. Z stacks were collected for 45 cells, 0.5 µm apart for a total thickness of 5 µm with approximately 4 frames per second for 90 seconds.

### Lysosome tracking quantification

Trackmate (Tinevez et al. 2017), an automated particle tracking software, was used to visualize individual lysosomes within CAD somas and neurites and calculate mean speed and velocity for each spot. The LoG detector was set to filter for spots at 0.5 microns in size with a quality threshold of 1.3. Thresholds and spot filtering were adjusted to exclude any particles detected outside the boundaries of the cell. The simple LAP tracker was then selected to remove any tracks that split or merge across the time course. Detected spots were filtered to remove gross outliers and only include spots that appear in at least 45 frames. Mean speed and total distance measurements were recorded for 5 spots in each compartment. Direction of transport was determined with the “TrackScheme” window and tracking particle displacement over the course of the imaging period.

### Comparison with neuronal RNA stability measurements

RNA halflife and subcellular abundance data were downloaded (Loedige et al. 2023). Halflife measurements were retrieved from the supplemental data of the referenced publication. Neurite enrichments were calculated using the salmon/tximport/DESeq2 approach described above. Overlaps with HNRNPA2B1 CLIP data were also performed as described above.

## AUTHOR CONTRIBUTIONS

The project was conceived by J.L. and J.M.T. Experiments were performed by J.L., K.F.V., and G.B. Data was analyzed by J.L., N.M., and J.M.T. Conceptual intellectual contributions were made by N.M., H.A.R., and J.K.M. Access to key equipment was provided by J.K.M. The manuscript was written by J.L., K.F.V, and J.M.T. We thank Lisa M. Wood of the J.K.M lab for microscope and live cell imaging data analysis training. Funding for this project was provided by H.A.R., J.K.M., and J.M.T.

## DATA AVAILABILITY

All high-throughput sequencing data associated with this manuscript can be found at Gene Expression Omnibus accession number GSE275212.

## SUPPLEMENTARY TABLES

**Table S1**. DESeq2 output comparing neurite enrichment values between HNRNPA2 knockout and rescue samples. Log2FoldChange represents the log2 difference in neurite enrichment between the samples (KO - rescue). Padj represents the corrected p-value for that difference in neurite enrichment.

**Table S2**. DESeq2 output comparing neurite enrichment values between HNRNPB1 knockout and rescue samples. Log2FoldChange represents the log2 difference in neurite enrichment between the samples (KO - rescue). Padj represents the corrected p-value for that difference in neurite enrichment.

**Table S3**. Neurite enrichments in rescue and knockout samples for WT and RGG mutant HNRNPA2B1 transgenes. Log2foldchange represents the neurite enrichment of the RNA (neurite / soma). Padj represents the corrected p-value for the differences in RNA abundances between soma and neurite.

## SUPPLEMENTARY FIGURES

**Figure S1.**
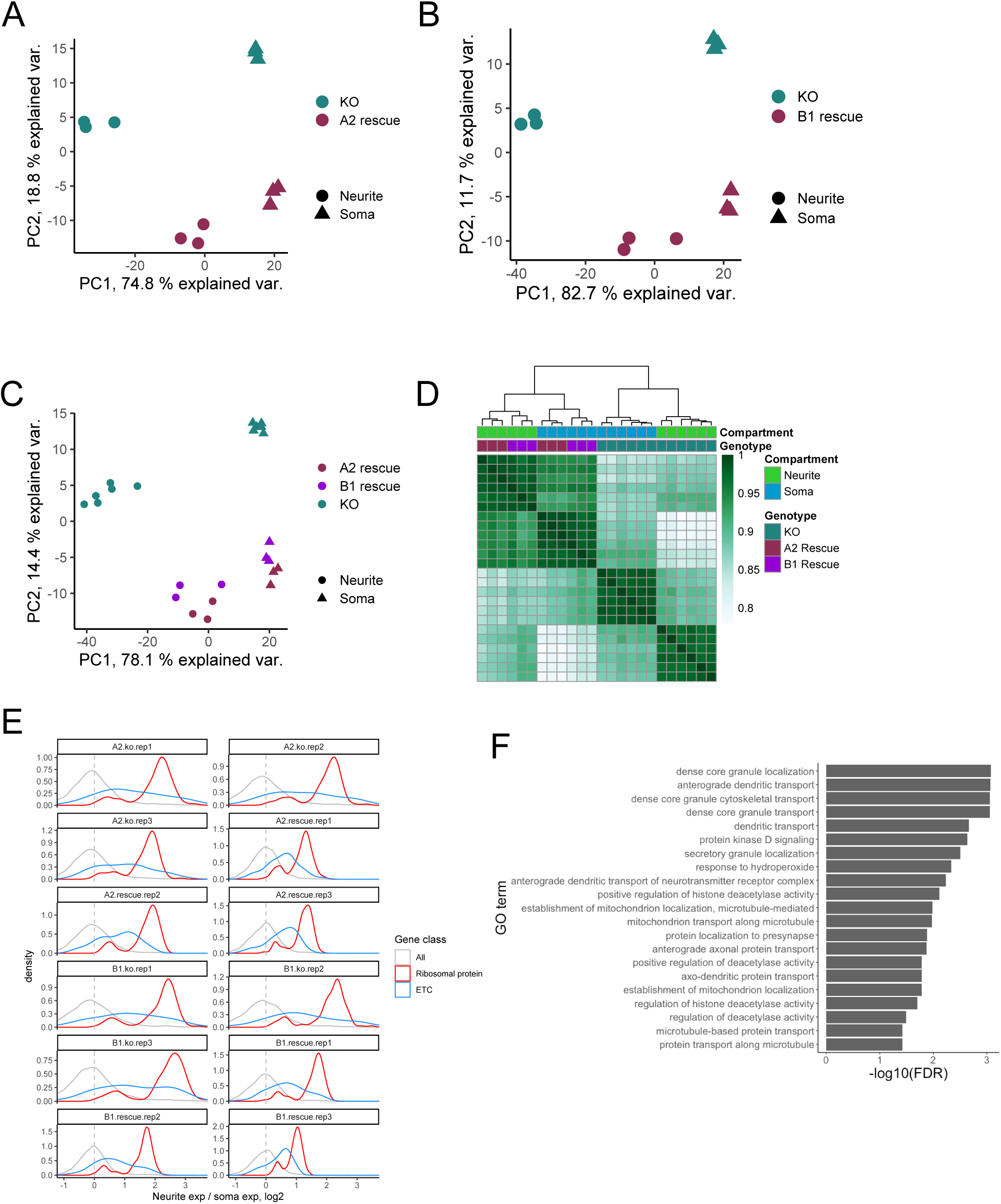
(A) Principal components analysis of gene expression levels from the soma and neurite of HNRNPA2B1 knockout and HNRNPA2 rescue cells. (B) Principal components analysis of gene expression levels from the soma and neurite of HNRNPA2B1 knockout and HNRNPB1 rescue cells. (C) Combined analysis of HNRNPA2 and HNRNPB1 rescue samples. (D) Hierarchical clustering of gene expression levels from HNRNPA2B1 knockout, HNRNPA2 rescue, and HNRNPB1 rescue soma and neurite samples. (E) Neurite enrichment levels for all RNAs (gray), those encoding ribosomal proteins (red), and those encoding components of the electron transport chain (blue). (F) Gene ontology terms enriched among the HNRNPA2B1-target RNAs, i.e. those that were mislocalized upon loss of HNRNPA2B1.

**Figure S2.**
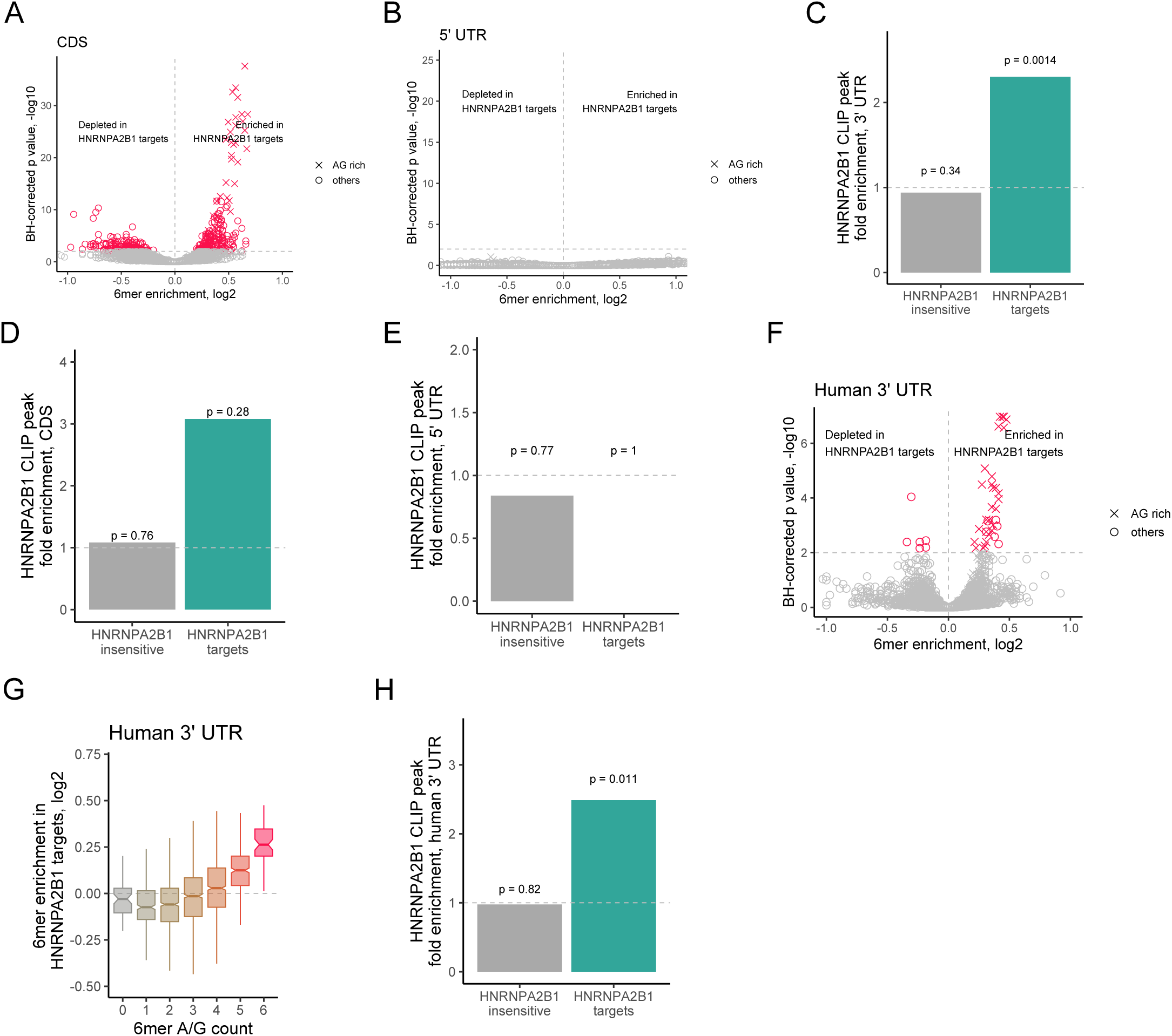
(A) Enrichment of 6mers in the coding sequences of HNRNPA2B1-targets compared to HNRNPA2B1-insensitive RNAs. 6mers comprised solely of adenosine and guanosine are marked with X. (B) As in A, but comparing the 5′ UTRs of HNRNPA2B1-targets and HNRNPA2B1-insensitive RNAs. (C) HNRNPA2B1 CLIP peak enrichment in the 3′ UTRs of HNRNPA2B1-target and HNRNPA2B1-insensitive RNAs. The dotted gray line represents frequency of HNRNPA2B1 CLIP peaks in the 3′ UTRs of all genes, normalized to 1. P values were calculated using a binomial test. (D) As in C, but looking at the coding sequences of the indicated RNAs. (E) As in C, but looking at the 5′ UTRs of the indicated RNAs. (F) As in A, but comparing the 3′ UTR sequences of the human orthologs of the HNRNPA2B1-target and HNRNPA2B1-insensitive RNAs. (G) Distribution of enrichments of 6mers with varying numbers of A/G residues in the 3′ UTRs of the human orthologs of the HNRNPA2B1-target and HNRNPA2B1-insensitive RNAs. (H) As in C, but comparing the 3′ UTRs of the human orthologs of the HNRNPA2B1-target and HNRNPA2B1-insensitive RNAs.

